# Experience-Dependent Plasticity in Nucleus Laminaris of the Barn Owl

**DOI:** 10.1101/2023.02.02.526884

**Authors:** Catherine E. Carr, Tiffany Wang, Ira Kraemer, Grace Capshaw, Go Ashida, Christine Köppl, Richard Kempter, Paula T. Kuokkanen

## Abstract

Barn owls experience increasing interaural time differences (ITDs) during development, because their head width more than doubles in the month after hatching. We therefore hypothesized that their ITD detection circuit might be modified by experience. To test this, we raised owls with unilateral ear inserts that delayed and attenuated the acoustic signal, then measured the ITD representation in the brainstem nucleus laminaris (NL) when they were adult. The ITD circuit is composed of delay line inputs to coincidence detectors, and we predicted that plastic changes would lead to shorter delays in the axons from the manipulated ear, and complementary shifts in ITD representation on the two sides. In owls that received ear inserts starting around P14, the maps of ITD shifted in the predicted direction, but only on the ipsilateral side, and only in those tonotopic regions that had *not* experienced auditory stimulation prior to insertion. The contralateral map did not change. Experience-dependent plasticity of the ITD circuit occurs in NL, and our data suggest that ipsilateral and contralateral delays are independently regulated. Thus, altered auditory input during development leads to long-lasting changes in the representation of ITD.

**Significance Statement:** The early life of barn owls is marked by increasing sensitivity to sound, and by increasing ITDs. Their prolonged post-hatch development allowed us to examine the role of altered auditory experience on the development of ITD detection circuits. We raised owls with a unilateral ear insert and found that their maps of ITD were altered by experience, but only in those tonotopic regions that had *not* experienced auditory stimulation prior to insertion. Thus experience-induced plasticity allows the sound localization circuits to be customized to individual characteristics, such as the size of the head, and potentially to compensate for natural conductive hearing losses.

## Introduction

Barn owls (genus *Tyto*) are altricial, and blind and deaf when they hatch (Haresign and Moiseff, 1988; Köppl et al., 2005). Young barn owls begin to acquire auditory sensitivity around day 4 posthatch (P4), their eyes open by about P10, their heads grow to adult width by about P30, and their facial ruff and sensitivity to high-frequency sounds is complete by about P60 (Haresign and Moiseff, 1988; Rich and Carr, 1999a; Köppl et al., 2005; Campenhausen and Wagner, 2006; Kraemer et al., 2017).

Upon hearing onset, connections in brainstem nuclei are rapidly refined, and at this time appear most vulnerable to disruption by deafening and noise rearing (Kapfer et al., 2002; Werthat et al., 2008; Kandler et al., 2009; Sonntag et al., 2009; Popescu and Polley, 2010; Butler and Lomber, 2013; Polley et al., 2013; Clarkson et al., 2016; Babola et al., 2018; Persic et al., 2020). Unilateral hearing loss in particular has profound effects on binaural circuits (Moore et al., 1999; Moore and King, 2004; Sanes and Bao, 2009; Popescu and Polley, 2010; Persic et al., 2020). Recent reviews have outlined the effects of hearing loss on auditory brainstem plasticity (Sanes and Bao, 2009; Tzounopoulos and Kraus, 2009; Anderson et al., 2010; Keating and King, 2013; Friauf et al., 2015; Lauer et al., 2019; Persic et al., 2020; Rubio, 2020), and upon localization behavior in rats (Clements and Kelly, 1978), ferrets (Moore et al., 1999; Keating and King, 2013; Kumpik and King, 2019), gerbils (Maier et al., 2008) and barn owls (Knudsen, 2002; Bergan and Knudsen, 2007; Keuroghlian and Knudsen, 2007). Previous studies have shown clear refinement of inputs to the ITD sensitive neurons of the mammalian medial superior olive during development (Kapfer et al., 2002; Chang et al., 2003; Magnusson et al., 2005; Werthat et al., 2008; Winters and Golding, 2018).

In the present study, we examined the effects of unilateral conductive hearing loss on ITD coding in NL, the first binaural nucleus in the barn owl. These recordings were possible because NL contains large, well-ordered, reproducible maps of ITD (Carr et al., 2015), which we exploited to test the hypothesis that neural delays might be modified by experience (Cheng and Carr, 2007). The ITD maps are created by afferent delay lines whose responses sum in NL to create a binaural ITD-sensitive neurophonic (Kuokkanen et al., 2013). We reasoned that, if delays adjusted to temporally altered acoustic input, then the maps of ITD would be shifted compared to normal. To test our hypothesis, we first described the maps of ITD in adult owls (McColgan et al., 2014; Carr et al., 2015). We then fitted 8 young owls (from P21) with unilateral ear-canal inserts (acoustic filtering devices) designed to introduce a time delay to inputs from that ear (Gold and Knudsen, 1999; Köppl et al., 2012), and later compared their ITD maps with those of normal adult owls. The manipulated owls did not show altered maps of ITD (Köppl et al., 2012). Although abnormal auditory experience starting around P25 induces adjustments in tuning for binaural localization cues in owl midbrain (Gold and Knudsen, 1999, 2000a, 2000b), we reasoned that P20 might be past a critical period for adjustment of delays in the brainstem NL. We therefore raised 6 owls from about P14 with unilateral earmold plugs, and report on the results of both studies here.

## Methods

All experiments were carried out on barn owls of both sexes. Recordings from American barn owls (*Tyto furcata;* previously classified as *Tyto alba pratincola*) in Maryland conformed to procedures in the NIH Guidelines for Animal Research and were approved by the Animal Care and Use Committee of the Universities of Maryland. Experiments in Oldenburg were carried out on European barn owls *(Tyto alba)* following protocols and procedures approved by the authorities of Lower Saxony, Germany (permit No. AZ 33.9-42502-04-11/0337).

### Acoustic filtering devices and earmold plugs

In our first experiments (Köppl et al. 2012), eight owls (2 *T. furcata* and 6 *T. alba*) were raised with acoustic filtering devices (Gold and Knudsen, 1999). Two owls (*T. furcata*) were first binaurally occluded with dense foam rubber earplugs (E.A.R. Cabot) to limit auditory experience from 20 to 28 days of age (P20-P28), corresponding to the time that their ear canals were open but too narrow to accommodate the device. Around 29 days of age, the binaural foam plugs were removed, and the filtering device was sutured into the right ear canal. The other six owls (*T. alba*) were raised beginning with a modified acoustic filtering device that could fit a younger ear canal (5.5 - 7.5 mm diameter of the ear-canal part), which was then replaced with a full size (8.5 mm diameter) device when the ear canals were large enough (Table 1). One of the other six owls (*T. alba*) was raised with an early version of the filtering device that had a shorter and smaller diameter chamber; the acoustic properties of this device were not measured. All 8 owl chicks were raised to minimum age of 3 months wearing the devices (see Table 1).

**Table 1.** Ear plug manipulations

| Manipulation | Age at which plug/device first applied | Duration | Number |
| --- | --- | --- | --- |
| Foam plug followed by device | P20-28 (soft plug) P29 (device) | 3-6 months | 2 *T. furcata* |
| Small (5.5 - 7.5 mm diameter) device followed by large (8.5 mm diameter) device | P20-27 (small device) P26-33 (device) | 3-6 months | 6 *T. alba* |
| Earmold plug | P14-17 (replaced with growth) | 3 months | 6 *T. furcata* |

The foam earplugs and the acoustic filtering devices were sutured into place while the owls were anesthetized with isofluorane (1-2%). The acoustic filtering device was designed by Gold (Gold and Knudsen, 1999) to alter auditory experience, and custom-made from acetal delrin (Plastics SRT). A small, circular flange fit tightly to the inner walls of the ear canal, and the rest of the device was located just behind the preaural flap and in front of the facial ruff feathers. The device was designed to increase the path length of sound reaching the affected ear and to change the resonance properties of the ear canal while still providing a low-impedance pathway to the tympanic membrane.

Since the acoustic filtering device did not appear to affect the map of ITD in the NL, a further 6 owl chicks (*T. furcata*) were raised with unilateral impression earmold plugs (Gold Velvet, Oklahoma City) inserted into the left ear canal under 1% isofluorane anesthesia. These earmold plugs could be inserted at an earlier age (starting at P14) to limit auditory experience. At P14, the average diameter of the ear canal was about 1.2 mm, increasing to about 6.2 mm at P30 (Haresign and Moiseff, 1988). During this first month after hatching, the ear opening diameter increases fairly linearly (diameter *y* (in mm) at time *x* (in days) was y = 0.275 *x* - 2.05), and it was possible to insert an earmold plug after about P14. The 6 owls, plus 2 age matched controls, were hand-raised in a Brinsea TLC-4 incubator (N. Somerset, UK), with temperature adjusted every few days as their thermoregulation ability increased (Rich and Carr, 1999b). The earmold plugs were examined daily and replaced every few days as the ear canals grew until the owls were about 3 months old. The plugs were then removed and the owls were returned to the aviary to reach adulthood (> 6 months), after which their maps of ITD were measured. Block of the middle ear was not possible because the large interaural canal that connects the two ears (Gans et al., 2012; Kettler et al., 2016).

### Auditory Brainstem Responses (ABRs)

To determine if plug rearing led to long-lasting changes in sensitivity, we used auditory brainstem responses (ABRs) to measure sensitivity in free field in 4 adult owls that had been raised with an earmold insert, and in 4 age matched controls (Fig. 1A). The plugs were removed prior to ABR recordings. For comparison with previous measurements, we used the same protocol and equipment as in (Kraemer et al., 2017). Owls were anesthetized by intramuscular injection of xylazine and ketamine. Anesthesia was induced by intramuscular injections of 16 mg/kg/hr ketamine hydrochloride (“Ketavet”, Phoenix, St. Joseph, MO) plus 3 mg/kg/hr xylazine (“Xyla-ject”, Phoenix). Supplemental doses of ketamine and xylazine were administered to maintain a suitable plane of anesthesia. Owls were placed in a large anechoic chamber (I.A.C.), wrapped in a heating blanket (38°C) with their head positioned 30 cm away from the speaker. Lidocaine was applied to the skin prior to inserting the electrodes subcutaneously.

**Figure 1.**
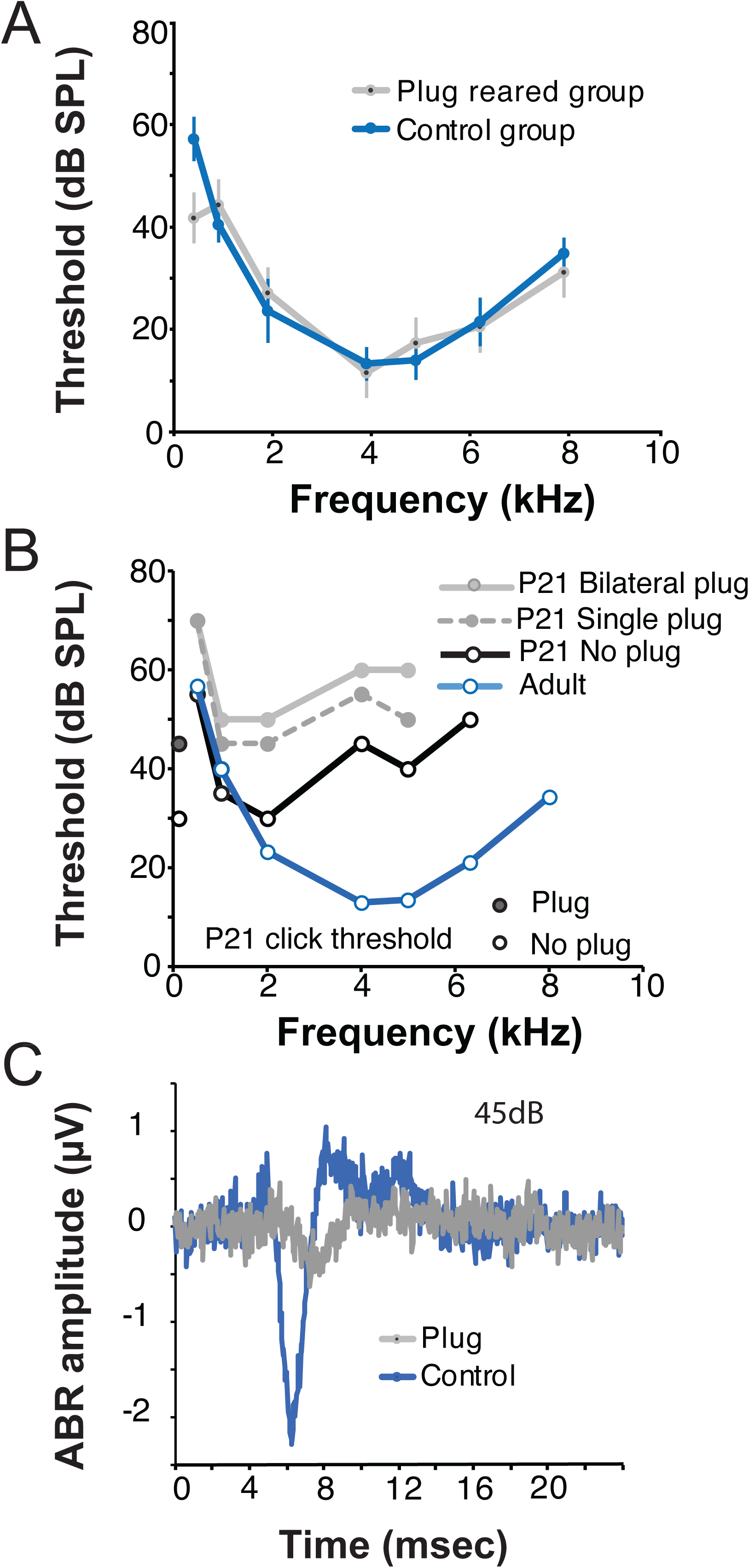
ABR evaluation of earmold plugs. A. Plug rearing does not affect overall hearing sensitivity, following removal of plug inserts. Average ABR audiograms for free-field tone-burst stimuli, for plug reared (inserts removed) and age matched control adults (n=8, 4 plugged and 4 control). Error bars show standard deviation in dB SPL. B. Earmold plugs induce reliable sound attenuation. ABR audiograms for P21 animals (mean of two), with no plug, a single plug or both ears plugged. Thresholds (for tone-bursts stimuli) were lowest with no plug insert, higher with a single plug, and highest with bilateral plugs. The adult audiogram is shown for comparison (Kraemer et al., 2017). Click thresholds for no plug or a single plug at P21 are shown at 0.1 kHz. C. Representative ABR traces from a single P21 animal before (blue) and after (grey) acute bilateral ear plug insertion. Traces were evoked by a free-field 45 dB click stimulus at t=0.

BioSigRP software (Tucker-Davis Technologies (TDT), Gainesville, Florida) was used to present stimuli and record from the three platinum subdermal needle electrodes: the negative electrode behind the ear facing directly towards the speaker, the ground electrode behind the opposite ear, and the active electrode directly on the midline vertex of the barn owl’s head. We used SigGenRP software (TDT) to create tone-burst stimuli, which were fed through an RP2.1 (TDT). The electrode signal was processed using a Medusa Digital Biological Amplifier System (RA4L Head stage and RA16PA PreAmp, RA16BA Medusa Base station). After data collection, the signals were notch filtered at 60 Hz, low-pass filtered at 3000 Hz, and high-pass filtered at 30 Hz using BioSigRP software. Thresholds for clicks and individual frequencies were established as the lowest stimulus level where any recognizable feature of the waveform was discernible, using visual inspection (Kraemer et al., 2017). Thresholds were defined as the decibel levels between the trace with a detectable peak and the trace without a detectable peak, so were either 5 dB or 2.5 dB SPL below the detectable peak. At least two thresholds (from 300 stimulus repetitions each) were determined for each frequency per trial. Thresholds were then averaged for each bird. ABR audiograms from owls raised with an earplug only differed from the age-matched control group at the 0.5 kHz point in the plug reared owls (Fig. 1A).

We also measured the effect of acute earmold plug insertion (unilateral and bilateral) in two P21 owls to determine the degree of attenuation caused by the earmold plugs around the time of insertion (Fig. 1B, C). We used the same ABR setup as before, with free-field stimulation. We measured the effect of both monaural and binaural earplug insertion and compared normal ABRs with ABRs recorded with bilateral and unilateral plugs (Fig. 1B). Above 2 kHz, mean bilateral attenuation for P21 animals was 16±3 dB, and for adult owls 17±4 dB (data not shown). An unbalanced one-way ANOVA showed significant differences in detection thresholds between normal and earplugged conditions for frequencies between 0.5-8 kHz in the adult owl (F_1,13_ =19.47, p<0.0007, not shown), and for frequencies between 0.5-6.3 kHz in the P21 chick (F_1,10_ =9.16, p<0.013, data in Fig. 1B, compare P21 bilateral and no plug).

### Surgery and stereotaxis and recording

To measure neurophonic potentials in NL, most animals were examined over 2 or 3 separate 8-hour experiments, spaced approximately a week apart, and designed to follow the protocols used in (Köppl et al., 2012; Carr et al., 2013, 2015). Anesthesia was induced by intramuscular injections of 16 mg/kg/hr ketamine hydrochloride ("Ketavet" Phoenix, St. Joseph, MO) plus 3 mg/kg/hr xylazine ("Xyla-ject", Phoenix). Supplementary doses of ketamine and xylazine were administered to maintain a suitable plane of anesthesia. Body temperature was measured by a cloacal probe and kept constant at 39°C by a feedback-controlled heating blanket wrapped around the owl’s body (Harvard Instruments, Braintree, MA). An electrocardiogram was used to monitor the plane of anesthesia.

The owl’s head was held in a controlled position by a custom-designed stereotaxic apparatus, using ear bars and a beak holder designed to position the head at 70° to the visual axis. Then a metal head plate and marker of a standardized zero point was permanently glued to the skull. After this, the ear bars and beak holder were removed, and the head held by the head plate alone. A craniotomy was made around the desired area relative to the zero point and a small hole made in the dura mater, taking care to avoid blood vessels. Each electrode was positioned at defined rostrocaudal and mediolateral locations, before being advanced into the brain. In some cases, the electrode was angled to facilitate access to the most medial regions of the brainstem.

Owls were placed on a vibration-insulated table within a sound-attenuating chamber (IAC, New York) that was closed during all recordings. Generally, commercial, Epoxylite coated tungsten electrodes (Frederick Haer Corporation, ME) were used, with impedances between 2 and 20 MΩ and generally 250 µm diameter shank; see Figure 8G in (Kuokkanen et al., 2010). A grounded silver chloride pellet, placed under the animal’s skin around the incision, served as the reference electrode. Extracellular electrode signals were amplified and filtered by a custom-built headstage and amplifier (mA 2000, Walsh Electronics, Pasadena, CA). Recordings were passed in parallel to an oscilloscope, a threshold discriminator (SD1, TDT) and an analogue-to-digital converter (DD1, TDT) connected to a personal computer via an optical interface (TDT). Analogue waveforms were saved for offline analysis.

### Acoustic stimulus generation and calibration

For neurophysiological recordings, acoustic stimuli were digitally generated by custom-written software ("Xdphys" written in Dr. M. Konishi’s lab at Caltech, CA) driving a signal-processing board (DSP2, TDT). After passing a digital-to-analogue converter (DD1, TDT) and an anti-aliasing filter (FT6-2, corner frequency 20 kHz, TDT), the signals were variably attenuated (PA4, TDT), impedance-matched (HB6, TDT) and attenuated by an additional fixed amount before being fed to commercial miniature earphones. Two separate channels of signals could be generated, passing through separate channels of associated hardware and driving two separate earphones (Yuin PK2, China). The earphones were housed in custom-built, calibrated from 0.1-10 kHz, closed sound systems inserted into the owl’s left and right ear canals, respectively. Sound pressure levels were calibrated individually at the start of each experiment, using built-in miniature microphones (Knowles EM3068, Ithasca, IL). The analogue stimulus waveforms were saved in most experiments.

Acoustic clicks, noises, and tone stimuli were digitally created with a sampling rate of 48.077 kHz (sampling interval: 20.8 µs). Clicks were digitally produced and then processed by the TDT-systems (Wagner et al., 2005, 2009). Several parameters of the click stimulus could be varied: intensity (maximally 0 dB attenuation, corresponding to 65 dB SPL spectrum level at 5 kHz (34 dB SPL overall level)), duration (1-4 samples equivalent to 20.8 - 83.2 µs), and polarity. A condensation click at 0 dB attenuation and 2 samples duration served as a reference. The click stimulus was repeated 128 times. Tone bursts of different frequencies (500 – 10,000 Hz range, 50 - 500 Hz steps) were presented monaurally and binaurally at levels from 20 - 60 dB SPL. Tone burst duration was 100 ms including 5-ms rise/fall times and with a constant starting phase. The noise had a spectrum between 0.1 and 13 kHz, 5- ms rise/fall times, and duration of 100 ms. The overall level of the noise varied between 20 - 60 dB SPL.

### Recording protocol

While lowering the electrode, noise bursts were presented as search stimuli. Once auditory neurophonic responses were discernable, tonal stimuli of varying frequencies were applied from both the ipsi- and contralateral side to judge the position of the electrode. To map best ITD within an isofrequency band, we made multiple stereotaxically controlled penetrations through the overlying cerebellum into NL. Apart from low-frequency NL, which has a lateral bend where frequencies from 500 - 2,000 Hz are represented (Takahashi and Konishi, 1988; Köppl, 1997, 2001), NL is sufficiently large and ordered for 3 - 4 serial penetrations along the same isofrequency slab within an 8-hour experimental period.

At a given recording site, the following protocol was tested to obtain a data set. To determine best frequency (BF) both ipsi- and contralaterally, average threshold crossings of the neurophonic potential in response to 3 - 5 repetitions of each stimulus were calculated within the stimulus window. This paradigm was chosen to derive iso-intensity frequency response curves for best frequencies between 2 - 7.5 kHz. Best ITD for each recording location was determined using two assumptions; first, that ITD tuning changed continuously with depth. Second, that peak responses to ITDs at or close to 0 µs would occur within a penetration. Best ITD were calculated from responses to either tone bursts presented binaurally at the estimated BF or to stimulation with noise bursts. Stimulus ITDs generally covered about two periods at the BF of a location. The tuning to ITD was judged by a Rayleigh test (p < 0.05). We recorded from both sides of the brain, with left ear leading ITDs described as negative, and right positive.

### Electrolytic lesions and histology

Lesions (10 µA, 3-10 s) were made at locations closest to 0 µs best ITD using a constant current device (CS3, Transkinetics, Canton, MA, or Lesion Making Device, Ugo Basile, Gemonio, Italy). After a survival time of 5 - 14 days, owls were perfused transcardially with saline, followed by 4% paraformaldehyde in phosphate buffer. Brains were blocked in the same stereotaxic apparatus as for *in vivo* recordings, marked on either left or right side, then cut in the transverse plane orthogonal to the electrode penetrations. Forty micron sections were mounted on gelatin-coated slides and stained with cresyl violet. A trained observer, blinded to condition, examined all sections at low magnification to identify electrode tracks and lesions, and then visualized and reconstructed every 3^rd^ section using a Neurolucida neuron tracing system (Microbrightfield Bioscience, St Albans, VT) that included an Olympus BX61 light microscope, a video camera (MBF-CX9000, Microbrightfield), a motorized stage controller, and a personal computer. Sections were traced and digitized using a 10X objective on the microscope, with the Z-position defined based on section thickness. Each section was aligned with the previous using rotation, reflection, or translation only to generate 3D reconstructions of NL. Lesions, lesion tracks, and NL borders were marked using the tracing function. The ruler function was used to measure distances from the lesion center to the borders of the nucleus. The distance of each lesion along its isofrequency band was measured from 3D reconstructions of NL. Isofrequency bands traverse NL and are composed of inputs from the nucleus magnocellularis with similar BF (Carr et al., 2015). Sections were not corrected for shrinkage (Kuokkanen et al., 2010, 2013) since measurements were normalized. Lesions varied in size, from 60 - 270 µm diameter (mean ± SD = 190 ± 64 µm, n = 61, Figures 3 and 6).

### Data analysis

A brief signal analysis was done during the experiment to determine frequency and ITD tuning. After the experiment, the responses were quantitatively analyzed with custom-written software (MATLAB; BEDS scripts from G. B. Christianson). Frequency and ITD tuning were quantified using the variance and the signal-to-noise ratio (SNR, (Kuokkanen et al., 2010)) of the ongoing neurophonic response; onset effects were excluded, and we averaged the variance and SNR over several trials. Only those recordings with both frequency and ITD tuning of the response variance (Kuokkanen et al., 2013; Carr et al., 2015) were accepted. The vector strength of the ITD tuning of the response variance was evaluated at each recording depth, and required to be > 0.02 and significant at p < 0.05 (Kuokkanen et al., 2013)

For analysis of ipsi- and contralateral BF tuning curves, the monaural iso-intensity frequency response curves were typically recorded multiple times within a penetration. Sites with weaker than the above ITD tuning were analysed if there was a strong ITD tuning of the cyclic mean amplitude (Kuokkanen et al., 2010, 2013) of vector strength > 0.1 and p < 0.005. Additionally, the monaural response curves needed to have maximum SNR (SNRmax) > 10 dB each. Apart from the direct comparison of the best frequencies (as defined in Kuokkanen et al. 2010 by the half-width at the half-height), we also cross-correlated the frequency response curves to ipsilateral and contralateral stimulation at each recording site to determine if there were a relationship between interaural mismatches in frequency tuning and ITD tuning. The BF response curves were linearly interpolated between stimulus frequencies to about 1-5 Hz resolution to ensure a smooth cross-correlation curve. The frequency lag at the maximum of the cross-correlation was used to describe the shift between the tuning curves.

### Statistics

We evaluated the effects of monaural occlusion on BF and best ITD using linear mixed-effects modeling in R statistical software (lme4 package). Linear mixed-effects models were chosen because they are robust to missing data and unbalanced designs. All earmold plugs had been placed to the left ear, which simplified analysis. Thus L_brain_ L_stimulus_ means recording from the side with plug (L_brain_), with ipsilateral stimulation (L_stimulus_), while L_brain_ R_stimulus_ means recording from the side with plug, with contralateral stimulation. The data were divided into low (≤5 kHz) and high (> 5 kHz) BF groups based on the stimulus frequency of the maximum SNR and stimulus frequency of the ITD tuning curve. The SNR_max_ of the neurophonic potential was incorporated as the response variable in each model, and recording side, BF-population (high or low BF frequency regions of NL), and their interaction (side x BF-population) were included as fixed effects. We incorporated subject ID as a random effect in each model to account for individual variation. Post-hoc analyses of significant fixed effects were performed by computing the estimated marginal means (i.e., least-square means) of linear mixed-effects models and examining pairwise comparisons among groups using the R package emmeans.

## Results

Since the development of stable ITD cues coincides with development of brainstem auditory circuits, we hypothesized that delays might be modified by experience (Fig. 2). We used the accessible map of ITDs in the NL to test this hypothesis in young owls raised with unilateral earplugs (Gold and Knudsen, 1999; Popescu and Polley, 2010; Anbuhl et al., 2017). Such inserts attenuate as well as delay the acoustic stimulus to the manipulated ear, in a frequency specific manner. It is important to note that ITD coding by the brainstem NL is based on phase-locked neural inputs whose preferred phase is basically invariant with sound level (review: (Heil and Peterson, 2015); barn owl: Köppl, 1997b; Wagner et al., 2005). Thus, although acoustic attenuation will additionally delay neural onset responses (Wagner et al., 2005), this effect is predicted to be negligible here, and the dominant effect of the ear inserts is assumed to be a unilateral, frequency dependent phase shift of the neural inputs to the developing NL on both sides of the brainstem. We predicted that, if compensation were to occur, the manipulated-ear inputs should become “faster”, i.e. show a decrease in delay when the plug is removed. If compensation were comprehensive, changes of up to 50 µs are to be expected, which corresponds to the typical acoustic delay effected by the ear inserts for frequencies up to 5-6 kHz (Knudsen et al., 1984; Gold and Knudsen, 1999). At still higher frequencies, the introduced acoustic delays are smaller.

**Figure 2.**
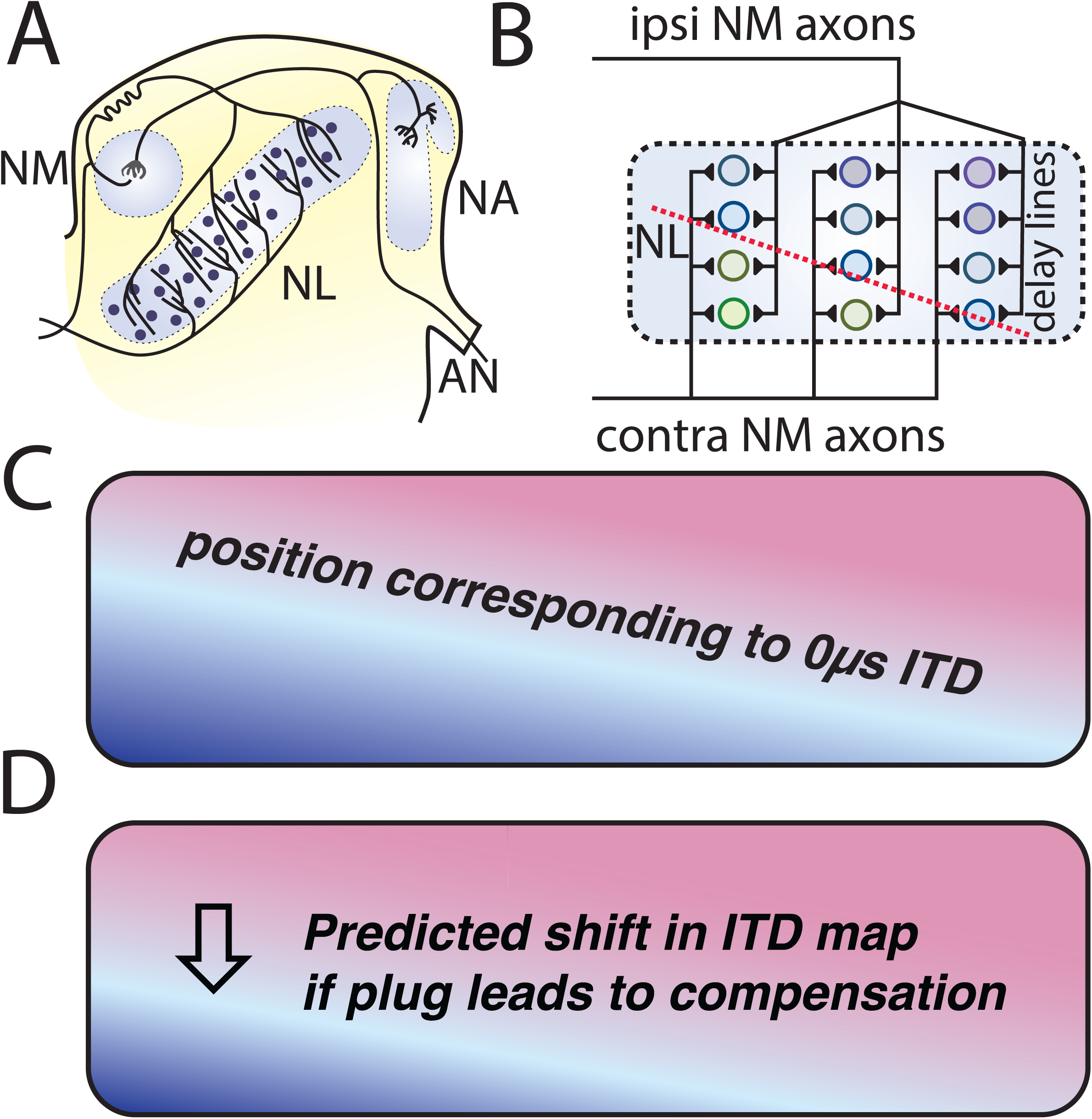
Predicted shift in ITD representation. A. Schematic connections from the auditory nerve (AN) to the first order nucleus magnocellularis (NM) and nucleus angularis (NA), and to the nucleus laminaris (NL), on right hand side (rhs) of the brainstem, from (Ashida and Carr, 2011). B. Schematic map of NL (rhs of the brainstem), with delay line axons from NM interdigitating in NL to create the maps of ITD. The dashed red line marks the position of 0 µs best ITD. Medial in the brainstem is to the left. C, D. Diagrams of rhs ITD maps in NL. Blue shading shows locations of best ITDs corresponding to sound sources in the ipsilateral hemifield (i.e., originating from the right side), red shading shows contralateral ITDs, and the pale diagonal band shows 0 µs or frontal ITD, from (Carr et al., 2015). C. Normal map, also predicted if there is no compensation for the altered auditory experience. D. Predicted shift in ITD representation after insert removal if there were compensation for ipsilateral plug insert rearing.

**Figure 3.**
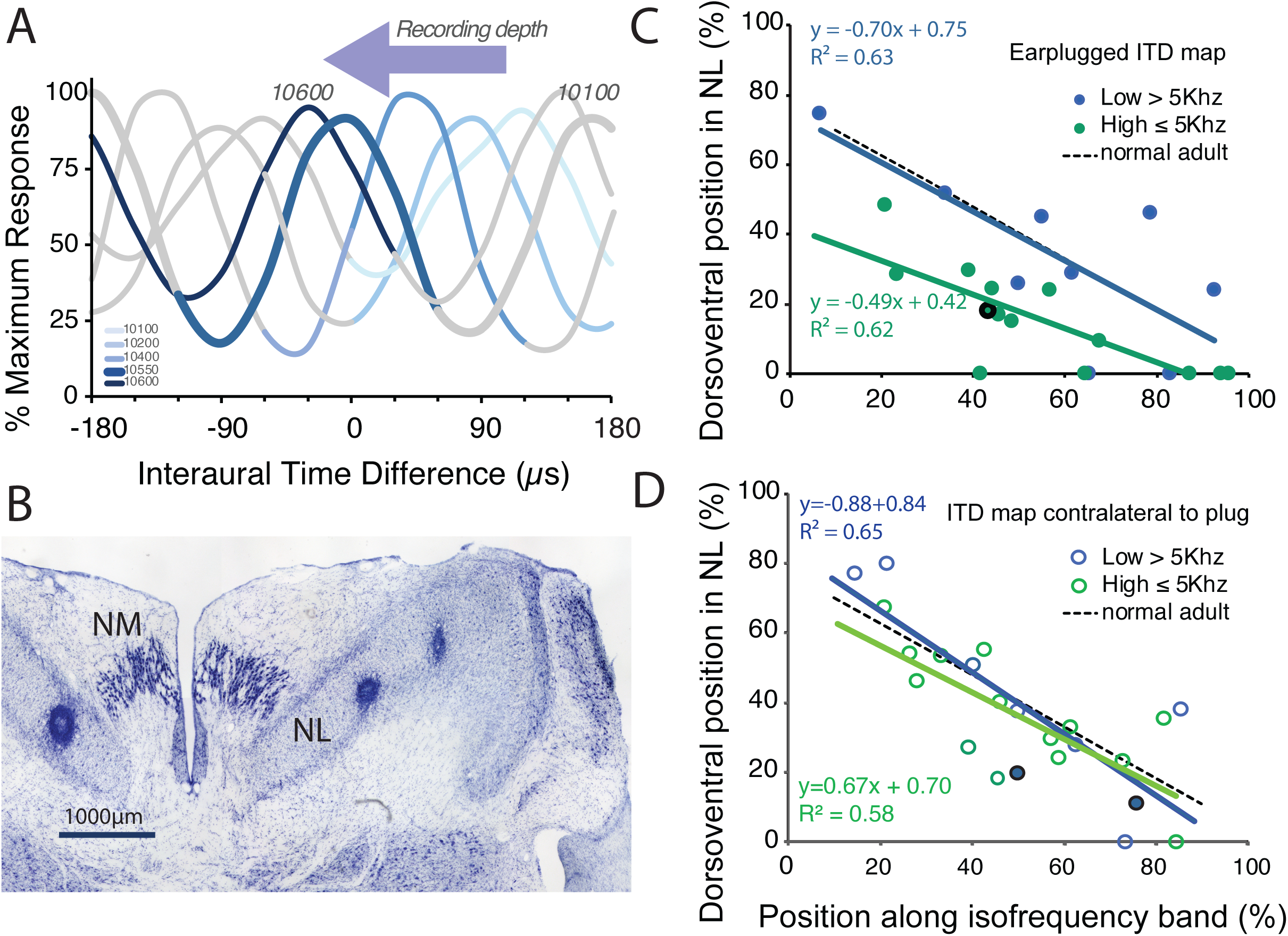
Maps of ITD in NL. A. Exemplar ITD sensitive recordings through left-hand side (lhs = plug side) NL. At each recording position (depth of electrode denoted in µm), the response shows local maxima at the best ITDs. There is a shift in best ITD from contralateral (positive best ITDs) to frontal ITDs (near zero best ITD) with increasing recording electrode depth (from dorsal to ventral). In this dorso-ventral series of recordings, with depth denoted by increasing shades of blue, lightest blue line shows most dorsal recording at 10,100 µm, thickest blue line has best ITD closest to 0 µs, and darkest blue line shows most ventral recording at 10,600 µm depth. Note that ITD tuning curves are largely periodic, and only one cycle of each ITD tuning curve is shown in blue (rest in grey). Neurophonic responses in NL were evoked by binaural tones (5.6 kHz) that varied in ITD, and ITD-sensitive responses were recorded only within NL. B. Nissl stained transverse 40 µm brain section at the level of central NL showing exemplar lesions from penetrations through lhs and rhs NL. The dorsal (at the top) and ventral (at the bottom) borders of NL are outlined by a glial border that surrounds NL, with the tracts made up of NM axons above and below the borders. Lesions on both sides in NL mark 0-µs best ITD : on the left, a lesion 43% along the tonotopic axis, and 18% up from the ventral edge of NL (black-ringed symbol in C) and on the right, two lesions, one at 50% along the tonotopic axis and 20% from the ventral edge, and the other at 76% of the tonotopic axis and 11% from the ventral edge, both black ringed symbols in D. Scale bar=1000 µm. C. Unilateral earmold plug-rearing is correlated with a compensatory shift in the ipsilateral ITD map, supporting the prediction in Figure 2D. Locations of lesions in NL (green dots) revealed a systematic shift in the mapping of frontal space, around 0 µs best ITD, with respect to the normal map (black dashed line, data from Carr et al., 2015), in NL ipsilateral to the earmold plug. Each data point marks the dorso-ventral and isofrequency axis coordinates of a 0 µs best ITD recording site. The dorso-ventral position of lesions marking 0 μs best ITD in NL decreased with increasing position along an isofrequency band (more lateral). Blue symbol colors denote recordings from regions of NL with best frequencies ≤ 5 kHz, and green symbols recordings > 5 kHz. Regression line colors match symbol color. Note C and D share a common x-axis. D. ITD maps in contralateral NL are robust to unilateral earmold plug-rearing. Locations of lesions in NL contralateral to the earmold plug show no systematic shift in the mapping of frontal space. Blue symbol color denotes recordings from regions of NL with best frequencies ≤ 5 kHz, and green symbols recordings > 5 kHz. Regression line colors match symbol color. Note C and D share a common x-axis.

We were able to test the hypothesis that delays might be modified by experience because we had previously characterized the normal ITD map (McColgan et al., 2014; Carr et al., 2015). In a typical recording through NL in an unmanipulated animal, an electrode penetration from dorsal to ventral in the brainstem reveals an ordered representation beginning with best ITDs to far contralateral stimuli. These best ITDs progress towards more frontal ITDs, then shift to a smaller representation of ipsilateral space, covering a total range of ∼200 µs ITD (Carr et al., 2015) (Fig. 2, 3A). The transition from contralateral to ipsilateral ITDs is associated with recordings of best ITDs of about 0 µs, and in our experiments was always marked by a small electrolytic lesion for later reconstruction (Fig. 3B). On both sides, we identified the 0 µs points as precisely as possible using online analyses (mean ± SD = -3.8 ± 5.5 µs, n=61) prior to making a lesion. The best ITDs were confirmed off-line using measures of signal-to-noise ratio (SNR (Kuokkanen et al., 2010)) and found to be close to the online measurements (mean ± SD = -5.7 ± 10.0 µs, mean difference 1.8 ± 7.3 µs, n=61). We then reconstructed the lesion locations to determine if there were changes in the map of ITDs (Fig. 3A, B).

NL is tonotopically organized, with low best frequencies mapped caudally, and progressively higher frequencies rostrally. A band across the mediolateral dimension of NL, oriented along a diagonal of about 45° to the midline, represents a single frequency and is referred to as an isofrequency band (Takahashi and Konishi, 1988; Carr and Konishi, 1990; Carr et al., 2013). For each bird, we made multiple penetrations within each isofrequency band to characterize the ITD maps both ipsilateral and contralateral to the brain side that had experienced the earmold plug (Fig. 2, 3A). In Carr et al. (2015) we modeled the normal ITD map (red dashed line in Fig. 2B), obtaining the regression shown as a black dashed line in Fig. 3C, D. These data enabled us to look for a shift in the map of ITD after raising an owl with an ear insert. When tested with the ear insert removed, we hypothesized that a “faster” ipsilateral input should cause the representation of frontal space, around 0 µs ITD, to be mapped significantly more ventrally within NL (Fig. 2D).

### Owls raised with a unilateral earmold plug had altered maps of ITD

We mapped ITDs in 6 owls raised with an earmold plug first inserted around P14-17. In the age-matched control owls, as described above, ITD maps show a steady shift in the position of 0 µs best ITD in the mediolateral dimension within any isofrequency slab (shown schematically in Fig. 2B, C). Thus, 0 µs best ITD is mapped dorsal in medial NL and ventral in lateral NL (Carr et al., 2015). This shift follows the regression described above (Figs. 3C and 3D, dashed black lines). In the owls raised with an earmold plug, we found a systematic ventral-ward shift in the location of the lesions made at 0 µs ITD ipsilateral to the earmold plug (Fig. 3C, solid green line). In the contralateral NL, the maps of ITD were not statistically different from the normal maps of ITD (Fig. 3D, solid green line), i.e., they shared similar slopes and y-intercepts to those of the control owls (Carr et al., 2015).

We thus had correctly predicted a ventral-ward shift in the map of ITD on the side with the plug (Fig. 2D). On the brainstem side contralateral to the earmold plug, we had erroneously predicted a decreased contralateral delay (mediated by the same NM neurons on the manipulated side), which would then cause an equal and opposite dorsal-ward shift in the maps of ITD. In other words, we hypothesized that the maps of ITD in the ipsilateral and contralateral NL would shift in opposite directions, but instead observed a shift confined to one side of the brainstem.

### Maps of ITD were not plastic after the onset of hearing

At first, we combined results from all iso-frequency bands because, in normal birds, the range of ITDs did not change with frequency, at least within the 3-8 kHz range examined (Carr et al., 2015). However, we were only able to insert the earmold plug around P14-18, or around the age when the owl chicks could already hear frequencies between 500 Hz - 5 kHz (Köppl and Nickel, 2007; Kraemer et al., 2017). We therefore separated the results by frequency, i.e., position along the tonotopic axis. The ITD maps from the 3-5 kHz regions of NL (Fig. 3C, solid blue line) were not different to the normal ITD map (Fig. 3C, black dashed line) while the ITD maps from 5-7.5 kHz were shifted ventral (Fig. 3C, solid green line). The slopes of each regression were not different, but the y-intercepts were, suggesting a ventral-ward shift of about 30%, corresponding to a shift in best ITD of approximately 50 µs. Since owls only begin to hear frequencies above 5 kHz after P16 (Köppl and Nickel, 2007; Kraemer et al., 2017), these results were consistent with the prediction that only a brief period of auditory experience was required for the neural delay lines in the ITD circuit to mature. In summary, the maps ipsilateral to the plug shifted in dependence upon the tonotopic region examined (Fig. 3C). Only ITD maps in the high best frequency (BF) region (> 5kHz) of NL showed plastic shifts.

Multiple critical periods have been documented at different levels of the auditory system (Polley et al., 2013; Persic et al., 2020). Early in development, before the emergence of hearing, waves of synchronous activity propagate through the developing auditory brainstem and are hypothesized to refine the tonotopic maps that define ITD and ILD circuits binaural circuits (Friauf and Lohmann, 1999; Jones et al., 2001; Kandler et al., 2009; Sonntag et al., 2009; Clause et al., 2014; Babola et al., 2018). Many synaptic and ionic mechanisms contribute to this observed refinement and can exhibit their own patterns of maturation (for reviews see (Trussell, 1999; Sanes, 2002; Kandler and Gillespie, 2005; Kubke and Carr, 2006; Tzounopoulos and Kraus, 2009; Klug et al., 2012; MacLeod and Carr, 2012; Ngodup et al., 2015; Kohrman et al., 2021)).

### Balance of ipsi- and contralateral inputs

It might be possible to create ITD shifts of the kind we observed through imbalanced inputs to the coincidence detectors from each side. We therefore compared both the BF and the ITD sensitivity of the neurophonic responses in low-BF (≤ 5kHz, unshifted maps) and high-BF (> 5 kHz, shifted map) regions of the NL ipsilateral to the plug with matching responses from the contralateral NL, and with measurements from age-matched controls (Fig. 4). Computational modeling and analysis of monaural neurophonic responses have shown that the neurophonic potential in NL is generated by hundreds of statistically independent sources, suggesting the afferent delay lines from the nucleus magnocellularis are its primary contributors (Kuokkanen et al., 2010, 2013, 2018). We therefore used the neurophonic to quantify the strength of the response from each ear, using the signal-to-noise ratio (SNR) of the monaural neurophonic responses to tones (Fig. 4B-E). These analyses helped to separate the role of location in brain (left NL, ipsilateral to the plug vs. the normal or right NL) from experience (recording from ipsi- and contralateral inputs within each NL, where one input originated from the plug side, and the other from the unmanipulated side).

**Figure 4.**
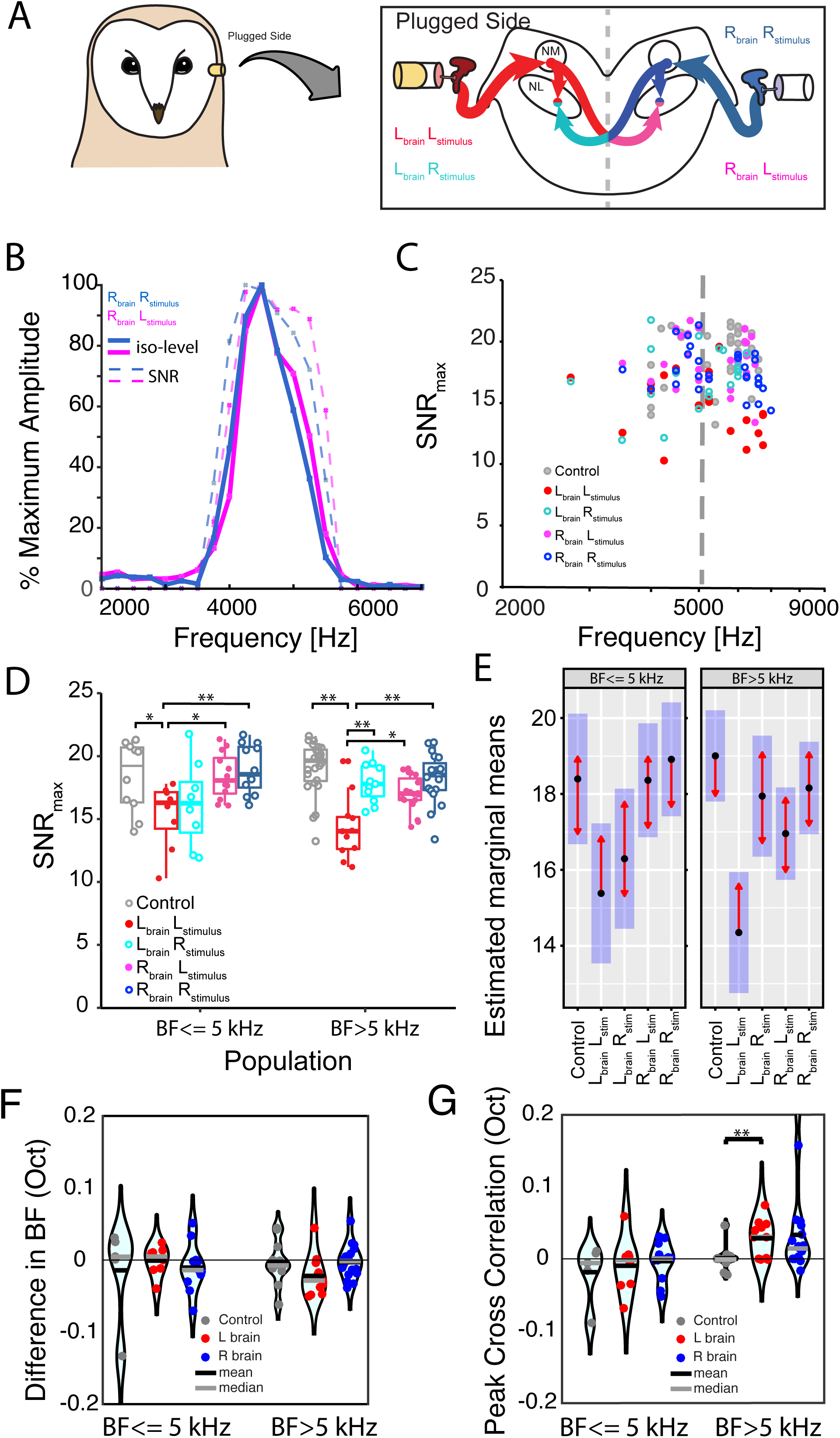
Plug effects on the strength of the inputs to NL. A. Diagram of earmold plug placement in the left ear, with schematic transverse section through the brainstem showing left (plug) ear inputs (in red color) to the left nucleus magnocellularis (NM). NM projects to both ipsilateral (red, L*_brain_* L*_stimulus_*) NL and to the contralateral (magenta, R*_brain_* L*_stimulus_*) NL. The right (no plug) ear inputs are shown in blue to the right NM. Right NM projects to ipsilateral (blue) NL, and to contralateral (cyan) NL. Color codes are maintained in the figure to facilitate comparisons. B. Exemplar iso-level neurophonic response curves (solid lines) recorded at a single location in right NL to stimulation of either the left ear (magenta, R*_brain_* L*_stimulus_*), or right ear (blue, R*_brain_* R*_stimulus_*). SNR (dashed lines) computed from each recording (SNR_max_ of 17.6 dB for L*_stimulus_*, 18.2 dB for R*_stimulus_*). C. SNR_max_ for all best frequency (BF) responses recorded on both left (plug, red and cyan) and right (no plug, magenta and blue) sides of the brain. Responses to stimuli on the same (=left) side as the earmold plug use filled symbols. Responses to stimulation of the right ear (no plug) use open symbols. Data on age-matched controls are shown in grey. The vertical dashed line divides the plot by frequency into low- (≤ 5 kHz) and high-frequency responses (> 5 kHz). D. Box plots of the same SNR_max_ data as in C (same color code), but separated for low (≤ 5 kHz) and high (>5 kHz) BF populations. Asterisks indicate significant pairwise differences in SNR_max_ for low- and high-BF regions of NL. For low-BF regions, SNR_max_ data for L_brain_L_stimulus_ (i.e., responses recorded on the brain side ipsilateral to the plugged (left) ear evoked by stimuli presented to the plugged ear) were significantly different from R_brain_R_stimulus_ (p<0.005) and R_brain_L_stimulus_ (p=0.027) responses and from age-matched controls (p<0.043). For high-BF regions, L_brain_L_stimulus_ recordings were significantly different from L_brain_R_stimulus_ (p=0.0012), R_brain_R_stimulus_ (p<0.0001), and R_brain_L_stimulus_ recordings (p=0.0178) as well as recordings from age-matched controls (p<0.0001). E. Comparison of estimated marginal means computed from linear mixed model analysis of all neurophonic SNR_max_ data from animals with earmold plugs and age-matched controls revealed that, for recordings with BFs ≤ 5 kHz, SNR_max_ of NL neurophonics recorded from the left side of the brain (ipsilateral to the plugged ear) and evoked by ipsilateral stimulation (L_brain_ L_stim_), were significantly different to those recorded on the right side of the brain (contralateral to the plugged ear) and evoked by ipsilateral and contralateral stimulation (R_brain_R_stim_ and R_brain_ L_stim_, respectively). For BFs > 5 kHz, SNR_max_ of L_brain_L_stim_ recordings were significantly different from all other groups. Confidence intervals for estimated marginal means are indicated by blue bars; pairwise comparisons are shown as red arrows with significant differences between groups indicated by non-overlapping arrows. F. Difference (in octaves) in BF for neurophonic responses to monaural ipsi- or contralateral stimulation (example in Fig 4B, solid lines), separately for NL on left (plug, red) side and right (no plug, blue) side of the brain, for all recordings, and age matched controls (grey). Positive difference corresponds to BF (L_stimulus_ ) > BF(control, R_stimulus_ ) independent of the recording side. No BF differences were significantly different from zero. G. To measure the difference in the frequency tuning, we correlated ipsi- and contralateral SNR tuning curves (example in Fig. 4B, dashed lines). The peaks of the correlation curve show a significant positive shift at the population level of high best frequency left brain (> 5 kHz, L_brain_ ) of BF (L_brain_L_stimulus_ ) - BF(L_brain_R_stimulus_ ) = 0.029 octaves ± SEM: 0.008 octaves, p = 0.004. No other BF differences were significantly different from zero.

We assessed the influence of rearing with a unilateral earmold plug on the SNR_max_ of neurophonic responses to monaural stimulation at BF using a linear mixed-effects model incorporating recording side (plug or no plug, each divided into recordings from ipsi- and contralateral inputs on both sides of the brainstem as described in the methods section), BF-population (above or below 5 kHz), and the interaction of side and BF-population as fixed factors, and subject ID as a random effect. The linear mixed model showed a strong (p<0.0001, Table 2) effect of the earmold plug on SNR_max_. Post-hoc pairwise comparisons between groups with a Tukey adjustment of p-values for multiple comparisons showed that ipsilateral inputs to NL on the same side as the earmold plug (L_brain_ L_stimulus_in Fig. 4A) were significantly weaker than the ipsilateral inputs to NL on the side contralateral to the earmold plug (R_brain_ R_stimulus_) and also weaker than the age-matched controls for both high- and low-BF populations (Fig. 4D, E). Although NM neurons project bilaterally, recordings from their axonal arbors showed reduced SNRs on the plug side (L_brain_ L_stimulus_; p<0.0001), and to a lesser extent on the unmanipulated side (R_brain_ L_stimulus_; P=0.0178) (Fig. 4E). In summary, the strength of inputs from the manipulated ear to coincidence detectors in NL on the plug side was reduced, with the strongest effect in the high BF regions of NL (Fig. 4C, D, E).

**Table 2.** Main effects of Side (plug or no plug) and BF Population (above or below 5 kHz) on SNR_max._

|  | Sum Sq | Mean Sq | NumDF | DenDF | F value | P |
| --- | --- | --- | --- | --- | --- | --- |
| Recording side | 191.314 | 47.829 | 4 | 117 | 10.516 | <0.0001** |
| BF-Population | 2.711 | 2.711 | 1 | 117 | 0.596 | 0.4417 |
| Recording side: BF-Population | 35.706 | 8.926 | 4 | 117 | 1.963 | 0.1047 |

**Table 3.**
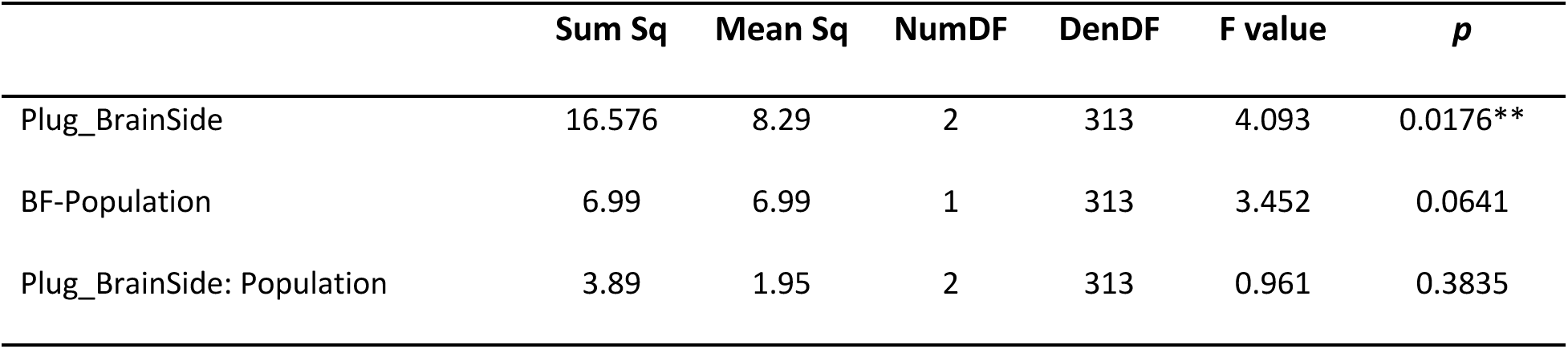
Main effects of Side (plug or no plug) and BF Population (above or below 5 kHz) on ITD

The stereausis hypothesis predicts that ITD tuning may be altered if ipsilateral and contralateral inputs originate from different cochlear positions, with different frequency tuning (Shamma et al., 1989; Pena et al., 2001; Carr et al., 2009; Plauška et al., 2017). We therefore investigated the relationship of interaural mismatches in frequency tuning and plug side (Fig. 4F). Ipsi- and contralateral inputs to NL were not always matched in BF, and sometimes the frequency mismatch could be large enough to affect ITD tuning. We found a significant difference between ipsi- and contralateral BF in high best frequency plug side groups (Fig. 4F, G, age-matched control owls). However, the observed BF mismatches, based on NM latencies Köppl (1997), only could account on average for -5.6 µs ITD shifts (±3.8 µs SD, range: -9.4 µs to 0.3 µs, Student’s 2 population T-test with respect to the high BF control population p = 0.0003, N = 9). Our data thus suggest that frequency tuning mismatches could contribute to the shifts in ITD tuning associated with rearing with a monaural earmold plug.

We also quantified the binaural ITD tuning at both plug and no-plug sides, using SNR_max_ to determine if there were a loss of synchrony associated with plug-rearing (Fig. 5E, F). The best ITD of each recording location was determined from the variance of the cyclic-mean signal (Kuokkanen et al., 2018), which corresponded to the circular-mean direction (mean phase) of the variance as a function of ITD (Fig. 5B, D). As before, we used a linear mixed model incorporating recording side (plug or no plug), BF- population (above or below 5 kHz), and the interaction of side and BF-population as fixed effects, and subject ID as a random effect. A statistical test of the model showed significant effects of side (p=0.0176); however, post-hoc pairwise comparisons found significant differences in SNR_max_ among recordings from the plugged and unplugged sides only for the high BF-population (p=0.0273, Fig. 5G). Thus, rearing with a unilateral earmold plug could be associated with a decrease in SNR.

**Figure 5.**
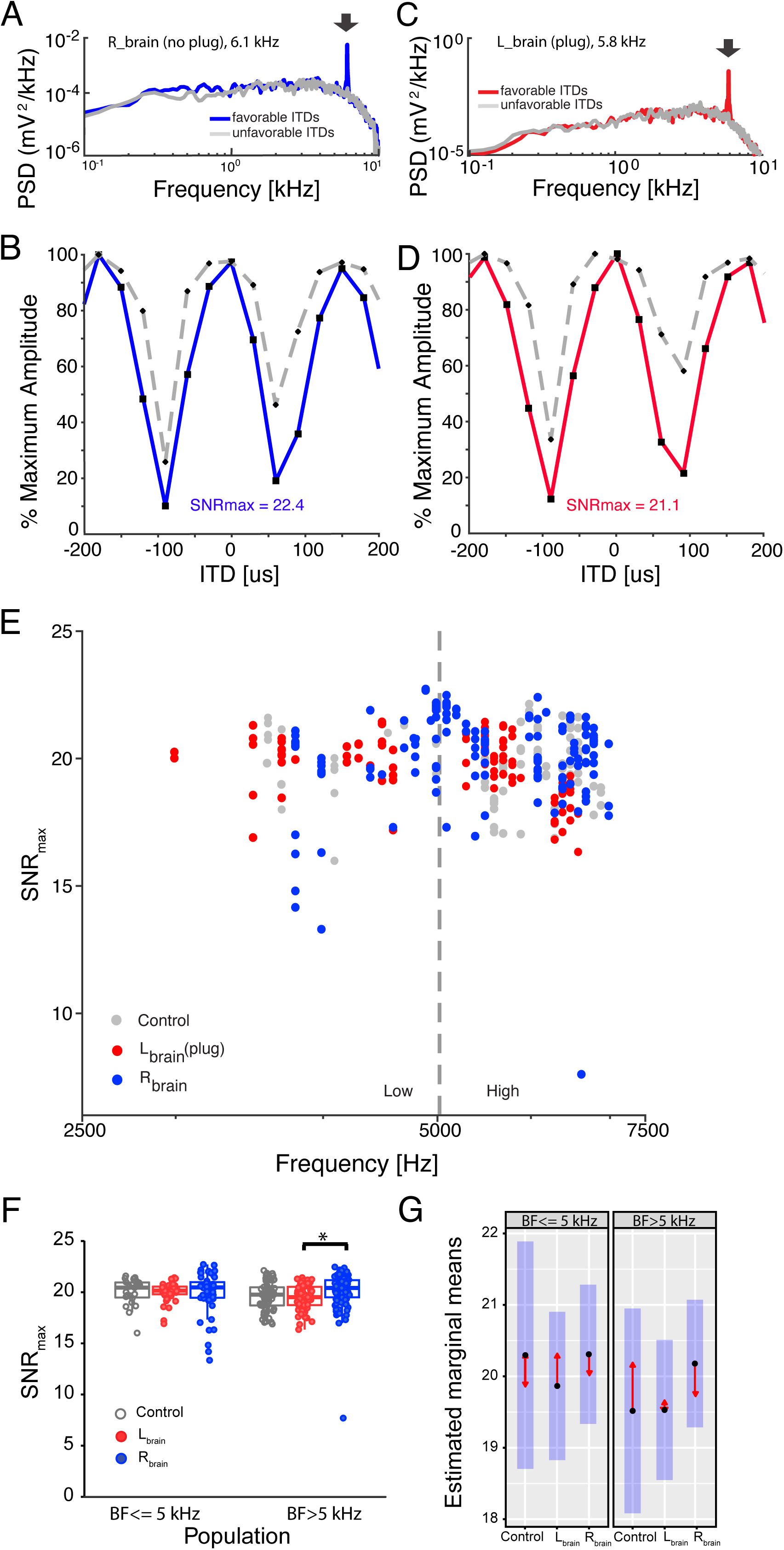
Plug effects on the ITD responses in NL on both sides of the brain. A. Mean power spectral density (PSD) of the binaurally driven neurophonic response in NL, for recording from the unmanipulated side, at favorable (blue curve) and unfavorable (grey curve) ITDs. Note the large “signal" peak (1.1·10^-2^ mV^2^/kHz) at the stimulus frequency for the favorable ITD (arrow at 6.1 kHz) and the average level around the peak (“noise”: 7·10^-5^ mV^2^/kHz ). For the PSD of the response to unfavorable ITD the signal peak height was 11·10^-5^ mV^2^/kHz and the noise level was 3·10^-5^ mV^2^/kHz. B. Exemplar neurophonic ITD curve (solid blue line) and SNR (grey dashed line), with BF tones as the stimulus, and responses recorded on the right (unplugged) side of the brain. The ITD curve was obtained from the variance of the sustained activity (onset excluded) across trials, where 100% corresponds to 0.79 mV^2^ in this example. The SNR is based on the signal and noise levels of the response PSDs to respective stimulus ITDs (examples in A), maximum SNR corresponds to 22.4 dB. Stimulation at 50 dB SPL; stimulus frequency 6.1 kHz. C. Mean PSD of the binaurally driven neurophonic response in NL, for recording from the side ipsilateral to the plug, at favorable (red curve) and unfavorable (grey curve) ITD. Note the large “signal" peak (0.78 mV^2^/kHz) at the stimulus frequency for the favorable ITD (arrow at 5.8 kHz) and the average level around the peak (“noise”: 6·10^-4^ mV^2^/kHz). For the PSD of the response to unfavorable ITD the signal peak height was 8·10^-4^ mV^2^/kHz and the noise level was 3·10^-4^ mV^2^/kHz. D. Exemplar neurophonic ITD curve (solid red line) and SNR (grey dashed line), with BF tones as the stimulus, and responses recorded on the left (plugged) side of the brain. 100% corresponds to 6.3 mV^2^ in this example (two example PSDs are shown in C), and the maximum SNR corresponds to 21.1 dB. Stimulation at 50 dB SPL; stimulus frequency 5.8 kHz. E. Maximum SNRs for all ITD-sensitive neurophonic recordings from animals raised with an earmold plug, separated according to brainstem side relative to the manipulated ear (L_brain_: ipsilateral to the plugged ear, red; and R_brain_: contralateral to the plugged ear, blue). Age-matched control measures of maximum SNR are also shown (grey). The vertical dashed line divides the plot into high (>5kHz) and low (≤5kHz) BF regions. F. Box and whisker plots of all maximum SNR values computed from ITD-sensitive neurophonic responses from animals raised with a monaural earmold plug. Data recorded from left (red, L_brain_) and right (blue, R_brain_), and age-matched controls (grey). For high BF regions, L_brain_ recordings were significantly different from R_brain_ (p=0.027, indicated on figure by asterisks). G. Comparison of estimated marginal means for the main effects of side and BF-population on SNR_max_, derived from linear mixed-effects modeling. Confidence intervals for estimated marginal means are represented as blue bars; red arrows indicate pairwise comparisons between groups where overlap of arrows indicates statistical non-significance.

### Maps of ITD were unaffected by rearing with a unilateral acoustic filtering device

In early experiments, unilateral acoustic filtering devices were implanted around P29, following foam earplugs at P20, to assess the role of experience in the development of sensitivity to ITDs in NL. The acoustic filtering device increased the path length of sound reaching the affected ear and changed the resonance properties of the ear canal while still providing a low-impedance pathway to the tympanic membrane (Köppl et al., 2012). A disadvantage of these filtering devices is that their large size did not permit early insertion in the ear canal. In contrast, the earmold plugs described above (Results in Figs. 3-5) could be placed in the ear canal as early as P14. When we used the unilateral acoustic filtering devices to assess the role of experience in the development of sensitivity to ITDs in 8 barn owl chicks, we found similar maps of ITD to the normal maps (Fig. 6B). The normal ITD map is characterized by a steady shift in the position of 0 µs best ITD (Carr et al., 2015), described by a regression of y = -0.74x + 0.78, R^2^=0.64 (Figure 6B, black line). In the owls raised with acoustic filtering devices, the maps of ITD were not statistically different to the normal owls (Fig. 6B, blue lines). The ITD maps did, however, appear to be less organized, i.e., the regressions explained less of the variance (Fig. 6)(Köppl et al., 2012). We hypothesize that brainstem and midbrain auditory circuits experienced different developmental windows for auditory plasticity.

**Figure 6.**
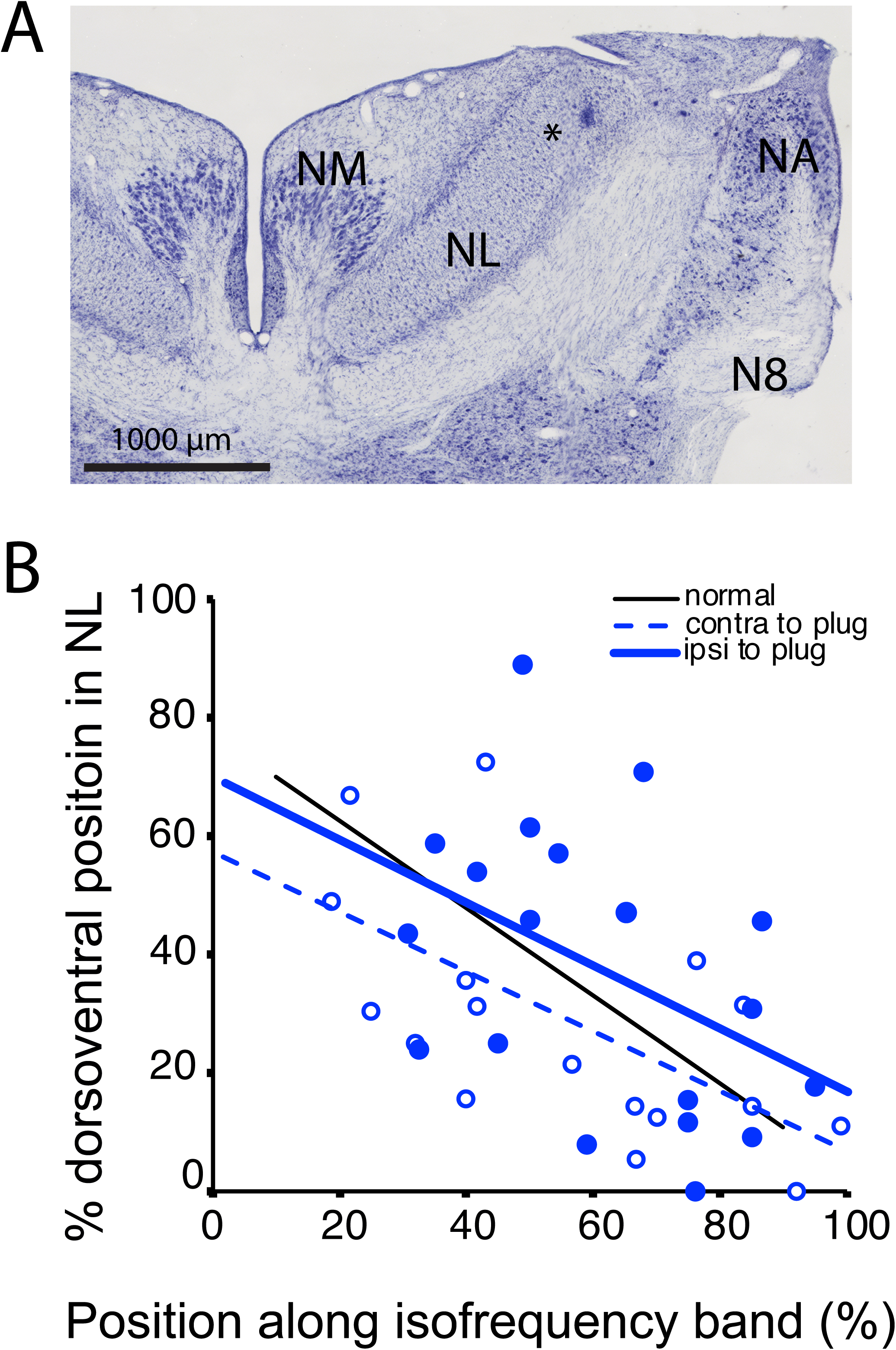
Maps of ITD in NL after device rearing with manipulations starting after P20. A. Example of a lesion used to identify the location of responses to 0 µs best ITD. Nissl-stained top half of rhs brainstem, showing NL, NM, nucleus angularis (NA), and the incoming auditory-nerve tract (N8) . Asterisk marks a lesion near the ventrolateral edge of NL (95% along isofrequency band). Scale bar = 1000 μm. B. Recorded ITD maps were not different to the normal maps in Carr et al. 2015 (solid line). ANCOVA with a z-test for slopes showed that the slopes were not different (p=0.3). A z-test for slopes yielded a p value of 0.64, thus the y-intercepts for the two ITD maps were not statistically different. Neither were they different to the normal map (p=0.70). Regression for line ipsilateral to the device y= -0.53x + 0.70, R² = 0.19. Regression contralateral to device y=-0.50x+0.57, R^2^=0.39.

## Discussion

Sensitivity to binaural delays can be modified when an ear is plugged prior to auditory experience (Knudsen et al., 1984; Popescu and Polley, 2010; Kumpik and King, 2019). We were able to examine the effects of these manipulations at the site of the first binaural comparisons because of the reproducible nature of the ITD maps in NL (Carr et al., 2015). There were long lasting changes in these maps, associated with a brief window for plasticity around the onset of hearing. Furthermore, changes were restricted to the side of the hearing loss, suggesting independent regulation of ipsilateral and contralateral circuits for detection of ITD.

### Effects of monaural occlusion

Young owls learn to associate ITDs with corresponding locations in space (Keller and Takahashi, 1996; Poganiatz and Wagner, 2001). If an ear is occluded during the first two months of life, owls were able to recover normal localization accuracy while plugged (Bergan 2007, Brainard 1995, Mogdans 1993). When the plug was removed, owls showed errors, but recovered localization (Knudsen et al., 1984). These behavioral changes are associated with significant remodeling of inferior colliculus circuits (Knudsen, 2002).

Knudsen’s experiments showed different sites and time constants for plasticity in the circuits for detecting both ITDs and interaural level differences (Mogdans and Knudsen, 1993; Gold and Knudsen, 2000b). Gold developed a phase-shifting acoustic filtering device to examine the plasticity in the barn owl ITD pathway. Rearing with this device inserted monaurally caused remodeling of the circuits within the inferior colliculus (Gold and Knudsen, 2000a, 2000b). When we raised young owls, using their paradigm, and with device insertion from P29, we also found no effect of altered auditory experience in the brainstem NL (Köppl et al., 2012), consistent with Gold’s findings. These results left unanswered the question of whether the brainstem representation of ITD could be modified by experience. Although the ear canals of younger owls were too small for the acoustic filtering device, we were able to induce a monaural conductive hearing loss beginning around P14-18. The extent of the shift in the ITD map was about 50 µs, consistent with the assumed delay imposed by these plugs (Knudsen et al., 1984). Thus, our data suggest different critical periods for binaural integration in brainstem and midbrain, as in other animal models of monaural occlusion (Popescu and Polley, 2010; Rosen et al., 2012; Polley et al., 2013; Mowery et al., 2015; Anbuhl et al., 2022).

In gerbils, monaural occlusion has well-defined effects upon the brainstem circuits for detection of ITD. Experiments with earplugs, monaural deafening and noise rearing in gerbils support a role for experience dependent segregation of inhibition to medial superior olive (MSO) cell bodies (Kapfer et al., 2002; Maier et al., 2008; Werthat et al., 2008; Sinclair et al., 2017). Recordings from the MSO’s target, the nucleus of the lateral lemniscus, related ITD sensitivity to the maturation of inhibitory inputs to the MSO (Seidl and Grothe, 2005). Thus, both *in vivo* recordings from the dorsal nucleus of the lateral lemniscus and studies of MSO support refinement in the days after hearing onset, similar to that observed for barn owl nucleus laminaris.

### Development of auditory sensitivity

To understand the effects of conductive hearing loss, it is important to know when animals can hear. In both birds and mammals, auditory responses typically begin with a restricted range of low to mid-frequencies (Romand, 1979; Brugge, 1983; Arjmand et al., 1988; Rubel et al., 1988; Sarro and Sanes, 2010). Rates of threshold maturation differ, depending upon where the animal falls along the altricial to precocial spectrum. Some mammals, such as guinea pigs (Dum, 1984) and humans (review in (Werner, 2007), are born with functioning auditory systems. Other mammals, such as gerbils (McFadden et al., 1996), ferrets (Moore, 1982) and cats (Walsh et al., 1986a, 1986b), are considered deaf at birth and altricial with respect to hearing (Pujol and Hilding, 1973; Beatty, 1998; Wess et al., 2017).

Birds show a low to high frequency pattern of auditory maturation (review in (Kubke and Carr, 2000, 2006)). Precocial birds respond to sound in the egg (Konishi, 1973; Gray and Rubel, 1985; Jones and Jones, 1995; Sato and Momose-Sato, 2003; Mann and Kelley, 2011; Rivera et al., 2018) with adult-like thresholds to low and middle frequencies by hatching (Saunders et al., 1973). Altricial bird hearing matures after hatching (Aleksandrov and Dmitrieva, 1992; Brittan-Powell and Dooling, 2004; Köppl and Nickel, 2007). When barn owl chicks hatch, they are insensitive to sound, and first show auditory responses to 1-2 kHz around P4-P7 (Köppl and Nickel, 2007; Kraemer et al., 2017). Sensitivity increases with age, with responses to 12 kHz appearing 2-3 months after hatching. Thus, owls experience a prolonged period of development that coincides with the maturation of the skull and ruff (Haresign and Moiseff, 1988; Hausmann et al., 2010).

The slow maturation of barn owl hearing allowed us to identify a critical period for auditory experience in the NL; we were able to insert an earmold plug when the young owls could hear frequencies up to 1-5 kHz (about 2 weeks after hatching) but were insensitive to higher frequency sounds (Köppl and Nickel, 2007; Kraemer et al., 2017). Since maps of ITD were only shifted in regions of NL with best frequencies above about 5 kHz, this suggests that experience is required for final maturation of the ITD circuit. These findings are consistent with the studies of the development of phase locking in the barn owl’s auditory nerve, in which phase locking emerges at about the same time as frequency tuning, over a prolonged period (Köppl, 2007). There are also strong parallels with results from the work on the gerbil brainstem, where activity dependent refinement occurs in the first days after hearing onset. Delays in maturation with white noise exposure suggest that not only activity, but patterned activity, is required for maturation of ITD tuning (Seidl and Grothe, 2005).

### Modification of ITD circuits

Our data suggest that the inputs to NL are plastic within a narrow time window around the onset of normal auditory experience, and furthermore, that the effects of monaural manipulation of acoustic experience appear predominantly ipsilateral. We had predicted that an acoustic delay caused by conductive hearing loss in one ear might be compensated for by a reduction in neural delay in the inputs to NL. We reasoned that these “faster” inputs could be detected after ear insert removal by a change in the ITD map, as a downward shift of the location of 0 µs best ITD in the ipsilateral map, and a matching upward shift on the contralateral side. The results were more complex than predicted. Only those parts of NL due to respond to frequencies that the owl could not hear at the time of plug insertion appeared to be plastic. Furthermore, only the ITD maps in the NL ipsilateral to the manipulated ear changed. One possible explanation for the prominence of ipsilateral changes is that ipsilateral conduction times may be more readily adjusted by shortening axonal projections, for example. In contrast, speeding up of the contralateral projection from the plugged side might be harder because these axons are already thicker to compensate for the longer distance (Seidl et al., 2010).

Identification of the mechanisms underlying modification of ITD circuits will require further study. Our analyses support limited compensation from misaligned frequency tuning or stereausis (Shamma et al., 1989; Pena et al., 2001; Carr et al., 2009; Plauška et al., 2017). According to the stereausis hypothesis, differences in wave propagation along the cochlea can provide delays necessary for coincidence detection if the ipsilateral and contralateral inputs originate from different cochlear positions, with different frequency tuning (Shamma et al., 1989). The stereausis model has not been supported in barn owls (Pena et al., 2001), alligators (Carr et al., 2009), and gerbils (Plauška et al., 2017), but could apply to circuit development. Models show that the necessary coherence in the signal arrival times could be attained during development, allowing learning to select connections with matching delays from amongst a broad distribution of axonal delays (Gerstner et al., 1996; Kempter et al., 2001).

The effect of side suggests the action of both or either of two mechanisms previously observed in gerbils; altered inhibition and changes in myelination. Gerbils show local and experience dependent changes in myelination and conduction velocity of inputs to the superior olive (Lehnert et al., 2014; Ford et al., 2015; Sinclair et al., 2017; Stange-Marten et al., 2017; Dutta et al., 2018), and similar mechanisms may apply to the developing barn owl circuit. Changes in inhibition could potentially compensate for the small phase delays measured in the barn owl NL, especially because inhibitory circuits allow for monaural effects; the superior olivary nuclei are the sole source of unilateral input to the nucleus laminaris (Carr 1989). In chicken, the avian superior olivary nuclei mutually inhibit each other (Burger et al., 2005). Thus, loud sounds in one ear should increase inhibition on the ipsilateral side, decrease inhibition on the contralateral side, and potentially alter response latency (Burger et al., 2011).

## Acknowledgements

We acknowledge Dr. E. Brittan-Powell for help with calibration and ABR recordings, and Dr. E. Smith for help with equipment and calibration. We thank Dr. Hermann Wagner and Sandra Brill for help measuring the neurophonic and Dr. Sahil Shah for establishing the lesion measurement paradigm. This research was sponsored by National Institute on Deafness and Other Communications Disorders (NIDCD) grant DC-000436 and DC-019341 (CEC). It was also supported by US-American Collaboration in Computational Neuroscience “Field Potentials in the Auditory System” as part of the National Science Foundation/NIH/French National Research Agency/German Ministry of Education and Research/United States-Israel Binational Science Foundation Collaborative Research in Computational Neuroscience Program, Research Grant 01GQ1505A. Research at the Carl von Ossietzky University of Oldenburg was supported in part by awards from the Hanse-WissenschaftKolleg, and the Alexander von Humboldt foundation.

## Literature cited

Aleksandrov LI, Dmitrieva LP (1992) Development of auditory sensitivity of altricial birds: absolute thresholds of the generation of evoked potentials. Neurosci Behav Physiology 22:132–137.

Anbuhl KL, Benichoux V, Greene NT, Brown AD, Tollin DJ (2017) Development of the head, pinnae, and acoustical cues to sound location in a precocial species, the guinea pig (Cavia porcellus). Hearing Research 356:35–50

Anbuhl KL, Yao JD, Hotz RA, Mowery TM, Sanes DH (2022) Auditory processing remains sensitive to environmental experience during adolescence in a rodent model. Nat Commun 13:2872.

Anderson S, Skoe E, Chandrasekaran B, Kraus N (2010) Neural timing is linked to speech perception in noise. J Neurosci 30:4922.

Arjmand E, Harris D, Dallos P (1988) Developmental changes in frequency mapping of the gerbil cochlea: Comparison of two cochlear locations. Hearing Res 32:93–96.

Ashida G, Carr CE (2011) Sound localization: Jeffress and beyond. Curr Opin Neurobiol 21:745– 751

Babola TA, Li S, Gribizis A, Lee BJ, Issa JB, Wang HC, Crair MC, Bergles DE (2018) Homeostatic Control of Spontaneous Activity in the Developing Auditory System. Neuron 99:511–524.e5

Beatty C Nell (1998) Structural Development of the Mammalian Auditory Pathways. In, pp 315– 413 Springer Handbook of Auditory Research. New York, NY: Springer New York.

Bergan JF, Knudsen EI (2007) Auditory Map Plasticity in Juvenile and Adult Owls. The Senses: A Comprehensive Reference, Vol 3, Audition:6.

Brainard MS, Knudsen EI (1995) Dynamics of visually guided auditory plasticity in the optic tectum of the barn owl. J Neurophysiol 73:595–614

Brenowitz S, Trussell LO (2001) Maturation of synaptic transmission at end-bulb synapses of the cochlear nucleus. J Neurosci 21:9487–9498

Brittan-Powell EF, Dooling RJ (2004) Development of auditory sensitivity in budgerigars (Melopsittacus undulatus). JASA 115:3092–3102

Brugge JF (1983) Development of Auditory and Vestibular Systems. Dev Auditory Syst:89–120.

Burger RM, Cramer KS, Pfeiffer JD, Rubel EW (2005) Avian superior olivary nucleus provides divergent inhibitory input to parallel auditory pathways. J Comp Neurol 481:6–18.

Burger RM, Fukui I, Ohmori H, Rubel EW (2011) Inhibition in the balance: binaurally coupled inhibitory feedback in sound localization circuitry. J Neurophysiol 106:4–14

Butler BE, Lomber SG (2013) Functional and structural changes throughout the auditory system following congenital and early-onset deafness: implications for hearing restoration. Frontiers Syst Neurosci 7:92.

Campenhausen M von, Wagner H (2006) Influence of the facial ruff on the sound-receiving characteristics of the barn owl’s ears. J Comp Physiol A 192:1073–1082

Caras ML, Sanes DH (2015) Sustained Perceptual Deficits from Transient Sensory Deprivation. J Neurosci 35:10831–10842

Carr C, Konishi M (1990) A circuit for detection of interaural time differences in the brain stem of the barn owl. J Neurosci 10:3227–3246

Carr CE, Shah S, Ashida G, McColgan T, Wagner H, Kuokkanen PT, Kempter R, Köppl C (2013) Maps of ITD in the nucleus laminaris of the barn owl. Adv Exp Med Biol 787:215–222

Carr CE, Shah S, McColgan T, Ashida G, Kuokkanen PT, Brill S, Kempter R, Wagner H (2015) Maps of interaural delay in the owl’s nucleus laminaris. J Neurophys 114:1862–1873

Carr CE, Soares D, Smolders JWT, Simon JZ (2009) Detection of interaural time differences in the alligator. J Neurosci 29:7978–7990

Chang EH, Kotak VC, Sanes DH (2003) Long-Term Depression of Synaptic Inhibition Is Expressed Postsynaptically in the Developing Auditory System. J Neurophysiol 90:1479–1488.

Cheng S-M, Carr CE (2007) Functional delay of myelination of auditory delay lines in the nucleus laminaris of the barn owl. Dev Neurobiol 67:1957–1974

Clarkson C, Antunes FM, Rubio ME (2016) Conductive Hearing Loss Has Long-Lasting Structural and Molecular Effects on Presynaptic and Postsynaptic Structures of Auditory Nerve Synapses in the Cochlear Nucleus. J Neurosci 36:10214–10227

Clause A, Kim G, Sonntag M, Weisz CJC, Vetter DE, Rubsamen R, Kandler K (2014) The precise temporal pattern of prehearing spontaneous activity is necessary for tonotopic map refinement. Neuron 82:822–835

Clements M, Kelly JB (1978) Auditory spatial responses of young guinea pigs (Cavia porcellus) during and after ear blocking. J Comp Physiol Psych 92:34–44.

Crins TTH, Rusu SI, Rodriguez-Contreras A, Borst JGG (2011) Developmental Changes in Short-Term Plasticity at the Rat Calyx of Held Synapse. J Neurosci 31:11706–11717

Dum N (1984) Postnatal Development of the Auditory Evoked Brainstem Potentials in the Guinea Pig. Acta Oto-laryngol 97:63–68]

Dutta DJ, Woo DH, Lee PR, Pajevic S, Bukalo O, Huffman WC, Wake H, Basser PJ, SheikhBahaei S, Lazarevic V, Smith JC, Fields RD (2018) Regulation of myelin structure and conduction velocity by perinodal astrocytes. Proc Natl Acad Sci USA 115:11832–11837

Dyke KBV, Lieberman R, Presacco A, Anderson S (2017) Development of Phase Locking and Frequency Representation in the Infant Frequency-Following Response. J Speech Lang Hear Res 60:2740–2751

Ford MC, Alexandrova O, Cossell L, Stange-Marten A, Sinclair J, Kopp-Scheinpflug C, Pecka M, Attwell D, Grothe B (2015) Tuning of Ranvier node and internode properties in myelinated axons to adjust action potential timing. Nat Commun 6:8073

Friauf E, Fischer AU, Fuhr MF (2015) Synaptic plasticity in the auditory system: a review. Cell Tissue Res 361:177–213.

Friauf E, Lohmann C (1999) Development of auditory brainstem circuitry. Activity-dependent and activity-independent processes. Cell Tissue Res 297:187–195

Gans E, Willis K, Bierman H, Carr C (2012) The interaural canal of the barn owl, Tyto alba. In: Integrative and Comparative Biology (ICB), pp e202–e356 Integrative and Comparative Biology

Gerstner W, Kempter R, Hemmen JL van, Wagner H (1996) A neuronal learning rule for sub-millisecond temporal coding. Nature 383:76–81

Gold JI, Knudsen EI (1999) Hearing impairment induces frequency-specific adjustments in auditory spatial tuning in the optic tectum of young owls. J Neurophys 82:2197–2209

Gold JI, Knudsen EI (2000a) A site of auditory experience-dependent plasticity in the neural representation of auditory space in the barn owl’s inferior colliculus. J Neurosci 20:3469– 3486

Gold JI, Knudsen EI (2000b) Abnormal auditory experience induces frequency-specific adjustments in unit tuning for binaural localization cues in the optic tectum of juvenile owls. J Neurosci 20:862–877

Gray L, Rubel EW (1985) Development of absolute thresholds in chickens. JASA 77:1162–1172

Haresign T, Moiseff A (1988) Early Growth and Development of the Common Barn-Owl’s Facial Ruff. Auk 105:699–705

Hausmann L, Campenhausen M von, Wagner H (2010) Properties of low-frequency head-related transfer functions in the barn owl (Tyto alba). J Comp Physiol A 196:601–612

Heil P, Peterson AJ (2015) Basic response properties of auditory nerve fibers: a review. Cell Tissue Res 361:129–158

Jones SM, Jones TA (1995) The tonotopic map in the embryonic chicken cochlea. Hearing Res 82:149–157

Jones TA, Jones SM, Paggett K (2001) Primordial rhythmic bursting in embryonic cochlear ganglion cells. J Neurosci 21:8129–8135

Kandler K, Clause A, Noh J (2009) Tonotopic reorganization of developing auditory brainstem circuits. Nature 12:711–717

Kandler K, Gillespie DC (2005) Developmental refinement of inhibitory sound-localization circuits. TINS 28:290–296.

Kapfer C, Seidl AH, Schweizer H, Grothe B (2002) Experience-dependent refinement of inhibitory inputs to auditory coincidence-detector neurons. Nat Neurosci 5:247–253.

Keating P, King AJ (2013) Developmental plasticity of spatial hearing following asymmetric hearing loss: context-dependent cue integration and its clinical implications. Front Syn Neurosci 7:123

Keller CH, Takahashi TT (1996) Responses to simulated echoes by neurons in the barn owl’s auditory space map. J Comp Physiol A 178:499–512 us&q=Responses+to+simulated+echoes+by+neurons+in+the+barn+owl’s+auditory+space+m ap&ie=UTF-8&oe=UTF-8.

Kempter R, Leibold C, Wagner H, Hemmen JL van (2001) Formation of temporal-feature maps by axonal propagation of synaptic learning. Proc Natl Acad Sci USA 98:4166.

Kettler L, Christensen-Dalsgaard J, Larsen ON, Wagner H (2016) Low frequency eardrum directionality in the barn owl induced by sound transmission through the interaural canal. Biol Cybern 110:333–343

Keuroghlian AS, Knudsen EI (2007) Adaptive auditory plasticity in developing and adult animals. Prog Neurobiol 82:109–121

King AJ, Parsons CH, Moore DR (2000) Plasticity in the neural coding of auditory space in the mammalian brain. Proc National Acad Sci 97:11821–11828

Klug A, Borst JGG, Carlson BA, Kopp-Scheinpflug C, Klyachko VA, Xu-Friedman MA (2012) How do short-term changes at synapses fine-tune information processing? J Neurosci 32:14058– 14063

Knudsen EI (2002) Instructed learning in the auditory localization pathway of the barn owl. Nature 417:322–328

Knudsen EI, Brainard M (1995) Creating a unified representation of visual and auditory space in the brain. Annu Rev Neurosci 18:19–43

Knudsen EI, Knudsen PF, Esterly SD (1984) A critical period for the recovery of sound localization accuracy following monaural occlusion in the barn owl. J Neurosci 4:1012–1020

Köppl C (1997) Frequency tuning and spontaneous activity in the auditory nerve and cochlear nucleus magnocellularis of the barn owl Tyto alba. J Neurophys 77:364–377

Köppl C (2001) Tonotopic projections of the auditory nerve to the cochlear nucleus angularis in the barn owl. JARO 2:41–53.

Köppl C (2007) Development of Phase Locking in the Barn Owl’s Auditory Nerve: The Emergence of Extreme Temporal Precision. ARO abstract.

Köppl C, Futterer E, Nieder B, Sistermann R, Wagner H (2005) Embryonic and posthatching development of the barn owl (Tyto alba): reference data for age determination. Dev Dyn 233:1248–1260

Köppl C, Nickel R (2007) Prolonged maturation of cochlear function in the barn owl after hatching. J Comp physiol 193:613–624

Kohrman DC, Borges BC, Cassinotti LR, Ji L, Corfas G (2021) Axon–glia interactions in the ascending auditory system. Dev Neurobiol 81:546–567.

Konishi M (1973) Development of auditory neuronal responses in avian embryos. Proc Natl Acad Sci USA 70:1795–1798.

Köppl C, Ashida G, Brill S, Kempter R, Wagner H, Carr C (2012) Experience-dependent plasticity in the nucleus laminaris of the barn owl. Front Behav Neurosci 6

Kraemer A, Baxter C, Hendrix A, Carr CE (2017) Development of auditory sensitivity in the barn owl. J Comp Physiol A 203:843–853

Kral A, Hubka P, Heid S, Tillein J (2013) Single-sided deafness leads to unilateral aural preference within an early sensitive period. Brain 136:180–193

Kubke MF, Carr C (2006) Development of the auditory centers responsible for sound localization. In: Springer Handbook of Auditory Research (Popper AN, Fay RR, eds), pp 179–237. Sound Source Localization.

Kubke MF, Carr CE (1998) Development of AMPA-selective glutamate receptors in the auditory brainstem of the barn owl. Microscopy Research and Technique 41:176–186

Kubke MF, Carr CE (2000) Development of the auditory brainstem of birds: comparison between barn owls and chickens. Hearing Research 147:1–20

Kumpik DP, King AJ (2019) A review of the effects of unilateral hearing loss on spatial hearing. Hearing Research 372:17–28

Kuokkanen PT, Ashida G, Carr CE, Wagner H, Kempter R (2013) Linear summation in the barn owl’s brainstem underlies responses to interaural time differences. J Neurophys 110:117– 130

Kuokkanen PT, Ashida G, Kraemer A, McColgan T, Funabiki K, Wagner H, Köppl C, Carr CE, Kempter R (2018) Contribution of action potentials to the extracellular field potential in the nucleus laminaris of barn owl. J Neurophys 119:1422–1436

Kuokkanen PT, Wagner H, Ashida G, Carr CE, Kempter R (2010) On the origin of the extracellular field potential in the nucleus laminaris of the barn owl (Tyto alba). J Neurophys 104:2274– 2290

Lauer AM, Dent ML, Sun W, Xu-Friedman MA (2019) Effects of non-traumatic noise and conductive hearing loss on auditory system function. Neuroscience 407:182–191.

Lehnert S, Ford MC, Alexandrova O, Hellmundt F, Felmy F, Grothe B, Leibold C (2014) Action potential generation in an anatomically constrained model of medial superior olive axons. J Neurosci 34:5370–5384

MacLeod KM, Carr CE (2012) Synaptic mechanisms of coincidence detection. In (Trussell LO, Popper AN, eds), pp 135–164 Synaptic Mechanisms in the Auditory System. Synaptic Mechanisms in the Auditory System.

Magnusson AK, Kapfer C, Grothe B, Koch U (2005) Maturation of glycinergic inhibition in the gerbil medial superior olive after hearing onset. J Physiology 568:497–512

Maier JK, Kindermann T, Grothe B, Klump GM (2008) Effects of omni-directional noise-exposure during hearing onset and age on auditory spatial resolution in the Mongolian gerbil (Meriones unguiculatus) — a behavioral approach. Brain Res 1220:47–57.

Mann ZF, Kelley MW (2011) Development of tonotopy in the auditory periphery. Hearing Research 276:2–15

McColgan T, Shah S, Köppl C, Carr C, Wagner H (2014) A functional circuit model of interaural time difference processing. J Neurophysiol 112:2850–

McFadden SL, Walsh EJ, McGee J (1996) Onset and development of auditory brainstem responses in the Mongolian gerbil (Meriones unguiculatus). Hearing Research 100:68–79

Miller GL, Knudsen EI (2003) Adaptive plasticity in the auditory thalamus of juvenile barn owls. J Neurosci 23:1059–1065

Mogdans J, Knudsen EI (1992) Adaptive adjustment of unit tuning to sound localization cues in response to monaural occlusion in developing owl optic tectum. J Neurosci 12:3473–3484

Mogdans J, Knudsen EI (1993) Early monaural occlusion alters the neural map of interaural level differences in the inferior colliculus of the barn owl. Brain Research 619:29–38

Moore DR (1982) Late onset of hearing in the ferret. Brain Res 253:309–311.

Moore DR (2009) Postnatal Development of the Mammalian Central Auditory System and the Neural Consequences of Auditory Deprivation. Acta Oto-laryngol 99:19–30.

Moore DR, Hine JE, Jiang ZD, Matsuda H, Parsons CH, King AJ (1999) Conductive Hearing Loss Produces a Reversible Binaural Hearing Impairment. J Neurosci 19:8704–8711

Moore DR, King AJ (2004) Plasticity of Binaural Systems. In: Plasticity of the Auditory System (Parks, Rubel, Popper EW, Fay, eds), pp 96–172 Springer Handbook of Auditory Research.

Mowery TM, Kotak VC, Sanes DH (2015) Transient Hearing Loss Within a Critical Period Causes Persistent Changes to Cellular Properties in Adult Auditory Cortex. Cerebral Cortex 25:2083– 2094

Ngodup T, Goetz JA, McGuire BC, Sun W, Lauer AM, Xu-Friedman MA (2015) Activity-dependent, homeostatic regulation of neurotransmitter release from auditory nerve fibers. Proc Natl Acad Sci USA:201420885

Parks TN, Rubel EW, Popper AN, Fay RR (2004) Plasticity of the auditory system. New York : Springer.

Pena JL, Viete S, Funabiki K, Saberi K, Konishi M (2001) Cochlear and neural delays for coincidence detection in owls. J Neurosci 21:9455–9459

Persic D, Thomas ME, Pelekanos V, Ryugo DK, Takesian AE, Krumbholz K, Pyott SJ (2020) Regulation of auditory plasticity during critical periods and following hearing loss. Hearing Res 397:107976.

Plauška A, Heijden M van der, Borst JGG (2017) A Test of the Stereausis Hypothesis for Sound Localization in Mammals. J Neurosci 37:7278–7289.

Poganiatz I, Wagner H (2001) Sound-localization experiments with barn owls in virtual space: influence of broadband interaural level different on head-turning behavior. J Comp Physiol A 187:225–233

Polley DB, Thompson JH, Guo W (2013) Brief hearing loss disrupts binaural integration during two early critical periods of auditory cortex development. Nat Commun 4:2547

Popescu MV, Polley DB (2010) Monaural Deprivation Disrupts Development of Binaural Selectivity in Auditory Midbrain and Cortex. Neuron 65:718–731

Pujol R, Hilding D (1973) Anatomy And Physiology Of The Onset Of Auditory Function. Acta Oto-laryngol 76:1–10

Rich V, Carr CE (1999a) Husbandry and captive rearing of barn owls. Poultry and Avian Biology Reviews 10:91–95.

Rich V, Carr CE (1999b) Husbandry and captive rearing of barn owls. Poul Av Bio Rev 10:91–95

Rivera M, Louder MIM, Kleindorfer S, Liu W, Hauber ME (2018) Avian prenatal auditory stimulation: progress and perspectives. Behav Ecol Sociobiol 72:112.

Romand R (1979) Development of auditory nerve activity in kittens. Brain Research 173:554– 556.

Rosen MJ, Sarro EC, Kelly JB, Sanes DH (2012) Diminished Behavioral and Neural Sensitivity to Sound Modulation Is Associated with Moderate Developmental Hearing Loss. Plos One 7:e41514.

Rubel EW, Parks TN, Edelman G, WE G, Cowan WM (1988) Organization and development of the avian brainstem auditory system. Auditory Function: Neurobiological Bases of Hearing.

Rubio ME (2020) Auditory brainstem development and plasticity. Curr Opin Physiology 18:7–10.

Sanes D (1993) The development of synaptic function and integration in the central auditory system. J Neurosci 13:2627–2637.

Sanes DH (2002) Right place at the right time. Nat Neurosci 5:187–188.

Sanes DH, Bao S (2009) Tuning up the developing auditory CNS. Curr Opin Neurobiol 19

Sarro EC, Sanes DH (2010) Prolonged maturation of auditory perception and learning in gerbils. Dev Neurobiol 70:636–648

Sato K, Momose-Sato Y (2003) Optical detection of developmental origin of synaptic function in the embryonic chick vestibulocochlear nuclei. J Neurophys 89:3215–3224

Saunders JC, Coles RB, Gates GR (1973) The development of auditory evoked responses in the cochlea and cochlear nuclei of the chick. Brain Research 63:59–74

Seidl AH, Grothe B (2005) Development of sound localization mechanisms in the mongolian gerbil is shaped by early acoustic experience. J Neurophys 94:1028–1036

Seidl AH, Rubel EW, Harris DM (2010) Mechanisms for adjusting interaural time differences to achieve binaural coincidence detection. J Neurosci 30:70–80

Shamma SA, Shen NM, P PG (1989) Stereausis: binaural processing without neural delays. JASA 86:989–1006

Sinclair JL, Fischl MJ, Alexandrova O, Heβ M, Grothe B, Leibold C, Kopp-Scheinpflug C (2017) Sound-Evoked Activity Influences Myelination of Brainstem Axons in the Trapezoid Body. J Neurosci 37:8239–8255

Sonntag M, Englitz B, Kopp-Scheinpflug C, Rubsamen R (2009) Early Postnatal Development of Spontaneous and Acoustically Evoked Discharge Activity of Principal Cells of the Medial Nucleus of the Trapezoid Body: An In Vivo Study in Mice. J Neurosci 29:9510–9520.

Stange-Marten A, Nabel AL, Sinclair JL, Fischl M, Alexandrova O, Wohlfrom H, Kopp-Scheinpflug C, Pecka M, Grothe B (2017) Input timing for spatial processing is precisely tuned via constant synaptic delays and myelination patterns in the auditory brainstem. Proc National Acad Sci 114:E4851–E4858.

Takahashi TT, Konishi M (1988) Projections of the cochlear nuclei and nucleus laminaris to the inferior colliculus of the barn owl. J Comp Neurol 274:190–211

Trussell LO (1999) Synaptic mechanisms for coding timing in auditory neurons. Annu Rev Physiol 61:477–496.

Tucci DL, Rubel EW (1985) Afferent influences on brain stem auditory nuclei of the chicken: Effects of conductive and sensorineural hearing loss on N. Magnocellularis. J Comp Neurol 238:371–381

Tzounopoulos T, Kraus N (2009) Learning to Encode Timing: Mechanisms of Plasticity in the Auditory Brainstem. Neuron 62:463–469

Wagner H, Brill S, Kempter R, Carr CE (2005) Microsecond precision of phase delay in the auditory system of the barn owl. J Neurophys 94:1655–1658

Wagner H, Brill S, Kempter R, Carr CE (2009) Auditory responses in the barn owl’s nucleus laminaris to clicks: impulse response and signal analysis of neurophonic potential. J Neurophys 102:1227–1240

Walsh EJ, McGee J, Javel E (1986a) Development of auditory-evoked potentials in the cat. II. Wave latencies. JASA 79:725–744

Walsh EJ, McGee J, Javel E (1986b) Development of auditory-evoked potentials in the cat. III. Wave amplitudes. JASA 79:745–754

Werner L (2007) Human Auditory Development. The Senses: A Comprehensive Reference, Vol 3, Audition:24.

Werthat F, Alexandrova O, Grothe B, Koch U (2008) Experience-dependent refinement of the inhibitory axons projecting to the medial superior olive. Dev Neurobiol 68:1454–1462

Wess JM, Isaiah A, Watkins PV, Kanold PO (2017) Subplate neurons are the first cortical neurons to respond to sensory stimuli. Proc Natl Acad Sci USA 114:12602–12607 Available at: http://www.pnas.org/lookup/doi/10.1073/pnas.1710793114.

Winters BD, Golding NL (2018) Glycinergic Inhibitory Plasticity in Binaural Neurons Is Cumulative and Gated by Developmental Changes in Action Potential Backpropagation. Neuron 98:166–178.e2

